# Amyloid-β induced membrane damage instigates tunneling nanotubes by exploiting p21-activated kinase dependent actin remodulation

**DOI:** 10.1101/655340

**Authors:** Aysha Dilna, Deepak K.V, Nandini Damodaran, Claudia S. Kielkopf, Katarina Kagedal, Karin Ollinger, Sangeeta Nath

**Author notes:** Corresponding to be addressed to Sangeeta Nath. Equally contributed (Aysha Dilna, Deepak K.V, Nandini Damodaran).

## Abstract

Alzheimer’s disease (AD) pathology progresses gradually via anatomically connected brain regions. Earlier studies have shown that amyloid-β1-42 oligomers (oAβ) can be directly transferred between connected neurons. However, the mechanism of transfer is not fully revealed. We observed formation of oAβ induced tunneling nanotubes (TNTs), nanoscaled f-actin containing membrane conduit, in differentially differentiated SH-SY5Y neuronal models. Time-lapse images showed that TNTs propagate oligomers from one cell to another. Preceding the TNT-formation, we detected oAβ induced plasma membrane (PM) damage and calcium-dependent repair through lysosomal-exocytosis and significant membrane surface expansion, followed by massive endocytosis to re-establish the PM. Massive endocytosis was monitored by an influx of the membrane-impermeable dye TMA-DPH and PM damage was quantified by propidium iodide influx in the absence of calcium. The massive endocytosis eventually caused accumulation of internalized oAβ in Lamp1 positive multi vesicular bodies/lysosomes via the actin cytoskeleton remodulating p21-activated kinase1 (PAK1) dependent endocytic pathway. Three dimensional quantitative and qualitative confocal imaging, structured illumination superresolution microscopy (SIM) and flowcytometry data revealed that oAβ induces activated phospho-PAK1, which modulates the formation of long stretched f-actin extensions between cells. Moreover, formation of TNTs can be inhibited by preventing PAK1 dependent internalization of oAβ using small-molecule inhibitor IPA-3, a highly selective cell permeable auto-regulatory inhibitor of PAK1. The present study gives insight that the TNTs are probably instigated as a consequence of oAβ induced PM damage and repair process, followed by PAK1 dependent endocytosis and actin remodeling, probably to maintain cell surface expansion and/or membrane tension in equilibrium.

## Introduction

Neurodegenerative diseases are propagating disorders characterized by accumulation of misfolded proteins that form aggregates, plaque and eventually cause neurodegeneration. A common hallmark of neurodegenerative diseases is prion-like self-propagation and gradual pathology progression in a predetermined pattern to different parts of the brain ^1^. Several studies have shown that proteins involved in these diseases such as tau, Aβ, α-synuclein and huntingtin follow common patterns including misfolding, self-propagation and neuron-to- neuron transfer ^2, 3^. In a model of Alzheimer’s disease (AD), we have previously shown that spreading of AD pathology is due to direct transfer of amyloidogenic oligomers between connected neurons ^4^. Moreover, lysosomal stress due to gradual accumulation of toxic non-degradable oAβ enhances the cell-to-cell progression of pathology ^5^. The studies ^4, 5^ have provided a possible explanations of how intracellular soluble oAβ, reported as the potential initiator or driver of AD ^6, 7^, could develop gradual pathology by propagating between connected cells. However, the mechanism of direct neuron-to-neuron propagation of neurodegenerative aggregates is not yet revealed.

Recently, several studies have demonstrated TNTs to transfer neurodegenerative proteins, such as PrP^Sc^, α-synuclein, Aβ, tau, polyQ, from one cell-to-another ^8–11^. Several of these studies implicated links between TNTs and the endo-lysosomal pathway in cell-to-cell spreading ^12^. In addition, exosomes are investigated as means of cell-to-cell transfer of Aβ ^13, 14^. However, these studies could not explain the anatomically connected strict spatiotemporal pathology progression of AD. On the other hand, cell-to-cell transfers of both extracellular and intracellular monomers and protofibrils of Aβ1-42 via tunneling nanotubes (TNTs) are demonstrated in primary cultures of neurons and astrocytes ^8^. However, the molecular basis of TNTs formation remains underexplored.

TNTs are open-ended membrane nanostructures consisting of membrane actin protrusions between neighbouring cells. Correlative cryo-electron microscopy has recently demonstrated that TNTs are formed by 2-11 individual TNTs and their diameters vary between 145 to 700 nm ^15^. TNTs are transient structures that can stay intact from minutes to hours. Membrane protrusions like filopodium precede TNT formation and inhibition of actin polymerization attenuates their formation ^16^. TNT formation prevails in neuronal cells and primary neurons ^17^. Successive studies also showed TNT formation in different cell types, such as immune cells, fibroblast, epithelial cells, astrocytes, and neurons, as well as their implication in the spreading of pathology in neurodegenerative diseases, HIV, herpes simplex virus (HSV) infections and in cancer ^18–20^. A growing number of studies have revealed that vesicle transfer, recycling vesicles, lysosomes and molecules involved in membrane expansion play a role behind the formation of TNTs, the actin membrane protrusions ^12, 21^.

In this study, we show oAβ induced formation of TNTs and direct cell-to-cell propagation of oAβ between neighbouring cells via TNTs. Preceding the formation of TNTs, we detect oAβ induced PM damage and repair through lysosomal exocytosis, which is followed by massive endocytosis via the membrane cytoskeleton actin remodulating kinase PAK1 engaged endocytic pathway. Endocytosis of oAβ activates phospho-PAK1, which modulates long stretched f-actin and formation of TNTs. Moreover, formation of TNTs can be prevented by inhibiting PAK1 dependent endocytosis and actin remodulation. Altogether, these observations give new insights that sprouting of TNTs might be instigated as a consequence of oAβ induced PM damage and Ca^2+^ dependent PM repair through lysosomal exocytosis via PAK1 dependent actin remodelling.

## Material and Methods

### Preparation of soluble oAβ

Freshly made unlabelled oAβ and fluorescently labelled oAβ- TMR were prepared from lyophilized Aβ (Aβ1–42) and Aβ-TMR (Aβ1–42-5-tetramethyl rhodamin) suspended in 1,1,1,3,3,3-hexafluoro-2-propanol (AnaSpec). Lyophilized Aβ and Aβ-TMR were resuspended at a concentration of 5 mM in Me2SO and then diluted to a concentration of 100 μM in HEPES buffer, pH 7.4. The solution was immediately vortexed and sonicated for 2 min and then incubated at 4°C for 24 hours ^4, 5^. Oligomers were characterized before the experiments similarly as reported in our earlier papers ^4, 5^, by electron microscopy imaging using a Jeol 1230 transmission electron microscope equipped with an ORIUS SC 1000 CCD camera, together with SDS-PAGE, Native-PAGE western blots and size exclusion chromatography.

### Neuronal cells culture and differentiations

SH-SY5Y neuronal cells (ECACC; Sigma-Aldrich) were seeded on 10-mm glass-bottom Petri dishes (MatTek) at a concentration of 12,000 cells/cm^2^. Cells were partially differentiated with 10 μM retinoic acid (RA; Sigma-Aldrich) for 7 days. Pre-differentiated or partially differentiated SH-SY5Y cells were further differentiated for additional 10 days in 10-mm glass-bottom Petri dishes (MatTek) with brain-derived neurotrophic factor, neuregulin-1, nerve growth factor, and vitamin D3. After 10 days of differentiation on glass, the cells form long, branched neurites and several neurospecific markers, as described previously ^4, 22^.

### Cell culture and transfections

For actin, PM, Lamp1 and GFP-GPI, plasmid constructs were used. The plasmid mEGFP-lifeact-7 (Addgene # 54610) was a gift from Michael Davidson and Lamp1-mGFP (Addgene # 34831) was a gift from Esteban Dell’Angelica ^23^. CAAX-mCherry (Ampicillin resistant) original source is ^24^ and GFP-GPI construct previously used by others ^25, 26^. The competent DH5α strain *E.coli* cells were used to transform the bacterial cells to be used for isolation of the plasmids. Plasmid DNA isolation was carried out using the QIAGEN Plasmid Midi kit. Transfections of SH-SY5Y cells were done using jetPRIME transfection reagent (Polyplus) and also by using Lipofectamine 3000 (Invitrogen). The cells (30,000 per well) were seeded on glass coverslips placed in a 24 well plate in 0.5 mL culture media. The plasmid DNA (0.5 µg) was diluted in 50 µL of jetPRIME buffer and vortexed for 10 seconds, followed by mixing with 1.6 µL of jetPRIME reagent which was then added to the cells cultured at 70-90% confluency and incubated between 24-48 hours before taking images. The transfection using Lipofectamine 3000 (Invitrogen) were done using (0.5 µg) plasmid-DNA complex added to the cells with fresh Opti-MEM (reduced serum) media which are at 70-90% confluency. The mixture was left for two to three hours and then removed. Fresh DMEM media with serum was added and microscopic images were taken after 24 - 48 hours.

### oAβ internalization/uptake

To investigate the uptake mechanisms of oAβ in the used cell system, differentiated SH-SY5Y cells were pre-treated with inhibitors in growth medium for the indicated time and at the indicated concentration (Table 1). After washing with PBS, a final concentration of 0.5 μM oligomerised Aβ-TMR was added to the cells for 1.5 h. After removal of the reagents, the cells were kept in a growth medium for 30 min before flow cytometry analysis. For flow cytometry, the cells were washed twice with PBS and trypsinised. Cells of 2 wells were resuspended in 300 μl PBS and filtered through a 50 μm nylon-mesh filter (Partec). The cells were analysed on a FACS Aria III flow cytometer (BD Biosciences) using FACSDiva acquisition software. Each treatment was carried out and analysed at least in triplicates. The gating was set using control cells without fluorophore and cells treated with Aβ-TMR only. The percentage of cells containing Aβ-TMR was calculated and normalised tothe mean of Aβ-uptake without any inhibitor, resulting in a fold change, to make results from different experiments comparable. Kaluza Software (Beckman Coulter) was used for data analysis.

**Table 1.** Pre-treatment conditions for inhibitors

| Inhibitor | Concentration | Treatment protocol |
| --- | --- | --- |
| α-Bungaratoxin | 100 nM | kept during experiment |
| AP-5 ((2R)-amino-5-phosphonopentanoate) | 100 μM | kept during experiment |
| GYKI 52466 [4-(8-methyl-9H-1,3-dioxolo[4,5-h][2,3]benzodiazepin-5-yl)-benzenamine hydrochloride] | 100 μM | kept during experiment |
| DPA (desipramine) | 25 μM | 2 h before experiment |
| Bafilomycin A1 | 10 nM | 30 min before experiment |
| NH<sub>4</sub>Cl | 10 mM | 30 min before experiment, no washing step, kept during experiment |
| MDC (monodansylcadaverine) | 50 μM | 30 min before experiment |
| PAO (phenylarsine oxide) | 300 μM | 30 min before experiment |
| Dynasore | 80 μM | 30 min before experiment |
| IPA-3 (1,1-Disulfanediyldinaphthalen-2-ol) | 20 μM | 30 min before experiment |
| EIPA (5-ethylisopropylamiloride) | 100 μM | 30 min before experiment |

### Colocalization of internalized oAβ

To study if oAβ was internalized within LAMP1 positive vesicles, different concentrations of oAβ-TMR of 250 nM, 500 nM and 1µM were added into the cells transfected with Lamp1-mGFP for time periods of 15 min, 30 min, 1 hour and 2 hours. oAβ (500nM and 1µM) internalization into the GPI positive vesicles were studied by incubating the oligomers for 15 min, 30 min and 1 hour into the cells transfected with GFP-GPI. Then images were taken after bleaching extracellular GFP-GPI by adding 0.4% Trypan blue (Sigma Aldrich) for 15 min and fixing with 4% PFA (Paraformaldehyde, Sigma Aldrich) for 15 min at room temperature and mounting with DABCO. oAβ (500nM and 1µM) and dextran FITC (1mg/ml) were co-incubated with the SH-SY5Y cells for 15 min, 30 min and 1 h min at 37°C. Then images were captured for both live cells and cells fixed with 4% PFA. Cells were visualised using fluorescence microscopy and % of colocalization from Mander’s overlapping coefficient were analyzed using ImageJ plugin Coloc 2 (open source by NIH, USA).

### Ca^2+^ dependent plasma membrane repair assay by propidium iodide staining

Undifferentiated SH-SY5Y cells were incubated with 1 μM of oAβ for 1 h or 2 h at 37 °C in the presence and absence of 5 mM EGTA (without Ca^2+^) in DPBS buffer (PBS with Ca^2+^ and Mg^2+^ chloride). Then the cells were washed and stained with propidium iodide (PI, 5 μg/ml) for 5 min. Cells were washed again two times with PBS before applying 4% PFA as fixative for 20 min at 4 °C. The fixed cells were observed by fluorescence microscopy or trypsinized before flow cytometry analysis.

### Immunocytochemistry

Lysosomal exocytosis was verified by immunocytochemical staining of Lamp1 on the outer leaflet of the PM in unfixed cells, using an antibody directed to the luminal part of Lamp1 in undifferentiated SH-SY5Y neuroblastoma cells as described earlier ^27^, on unfixed cells. Cells were incubated for 15 and 30 min with 1 µM of oAβ in MEM media without FCS (fetal calf serum) supplement. Then, endocytosis of the cells was blocked with 5% BSA + 10% FCS in PBS for 5 min at 4 °C. Then cells were incubated with Lamp1anti-goat primary antibody (1:250, sc-8099, Santa Cruz Bio-technology; Santa Cruz, CA, USA; 2 h, 4 °C) in the blocking buffer for 45 min, followed by fixation of the cells in 4% PFA for 20 min at 4 °C before labelling with the secondary anti-goat antibody conjugated to Alexa Fluor 488 (1:400 for 30 min; Molecular Probes, Eugene, OR, USA). Next, the cells were mounted in ProLong Gold antifade reagent supplemented with 4′,6-diamidino-2-phenylindole (DAPI; Molecular Probes). Conventional immunocytochemical staining was done to quantify the activated PAK1 and actin in oAβ and IPA-3 treated cells using phospho-PAK1 (Thr423)/PAK2 (Thr402) antibody (Cell signalling #260; 1:150) and Phalloidin–Tetramethylrhodamine B isothiocyanate (Sigma P1951, 1:500). Anti-rabbit Alexa 405 and FITC (1:500) were used as secondary antibodies.

### Live cell imaging of membrane dynamics

oAβ (1 µM) was added concurrently with 1 μM of the membrane binding dye TMA-DPH (N,N,N-Trimethyl-4-(6-phenyl-1,3,5-hexatrien-1-yl) phenylammonium p-toluenesulfonate; molecular formula: C28H31NO3S) (10 mM of TMA-DPH stock solution was made by dissolving in methanol) and membrane dynamics of the live cells were followed by time-laps images using a confocal microscope. The excitation and emission maximum of TMA-DPH is 384 and 430 nm, with a considerable tail of excitation spectra at 405 nm. Therefore, time-lapse images were taken using a confocal-microscope by exciting the TMA-DPH dye using the 405 nm laser and sufficient emission light was collected using an opened pinhole. In this setup confocal microscope produces images similar to the widefield or epifluorescence images. Cultures were carried to the microscope one by one in 500 µl of 20 mM HEPES buffer of pH 7.4 maintaining the temperature at 37°C. oAβ (1 μM) was added concurrently with 1 μM of membrane binding dye TMA-DPH to HEK cells similarly as above and membrane dynamics of the live cells were studied by images taken by fluorescent microscope.

### Cell viability assay

Viability of differentiated cells were measured in triplicates using MTT reagent with the undifferentiated cells treated with oAβ (1μM) for 1 h with or without IPA-3 (20 μM) pretreated cells for 30 min at 37°C. Cells were incubated with MTT reagent for 2 hours at 37°C, then removed all the media and measured the DMSO solubilized formazan. Formazan (bright orange in color), the reduced MTT product produced by viable cells after 2 h of incubation at 37°C was dissolved in DMSO and measured, at 570 nm using a Victor Wallac (PerkinElmer) plate reader.

### Confocal and fluorescence microscopy to image TNTs

Formation of oAβ induced TNTs was observed in the cells treated with increasing concentrations of oAβ at different incubation times, using either confocal or fluorescence microscopes. The treatments were done by incubating the cells with oAβ in serum free medium and corresponding control cells were treated at the same conditions, to nullify the serum starvation induced TNTs. Differentiated, partially differentiated and undifferentiated cells on glass petri dishes were incubated with 100–500 nM oAβ–TMR for 3 h at 37°C in a 5% CO2 atmosphere. The cells were imaged after incubating with LysoTracker (green; Invitrogen) 50-250 nM for 5-10 min and after extensive PBS washing (two washes of 10 min each at 37°C with 5% CO2). Images were acquired using a Zeiss LSM laser scanning confocal microscope using 63X/1.4 NA or 40X/1.3 NA oil immersion plan-apochromatic objective (Carl Zeiss AG, Oberkochen, Germany). The time-laps image sequences of the live cells were taken at 37°C by capturing simultaneously differential interference contrast (DIC) and fluorescence modes. Fluorescence microscopy (IX73-Olympus) with 63X/1.3 NA and 100X1.4 NA was also used to do the experiments of SH-SY5Y cells transfected with Plenti-lifeact EGFP plasmid and CAAX-mCherry plasmid. To study the effect of IPA-3 on SH-SY5Y cells, the cells were first treated with IPA-3 (20 μM) for 30 minutes followed by oAβ (1 μM) for 1 - 3 hours respectively. The cells were then imaged under microscopes and numbers of TNTs were quantified and plotted in percentage by manual counting the TNTs with respect to the number of cells from each of the image frames.

### Flow-cytometry

Internalization of oAβ-TMR in the presence and absence of inhibitors were quantified using BD FACS Aria ^TM^ (BD Biosciences) and were analysed using BD FACS DIVA ^TM^ (BD Bioscienc es) flow cytometer. Immunocytochemical stained cells with anti PAK1 and actin-phalloidin were fixed, trypsinised and suspended in PBS and quantified using BD LSR II (BD Biosciences) flow cytometer. Cells treated with propidium iodide (PI) were fixed, trypsinized and filtered using CellTrics 30 μm filters (Sysmex). Then re-suspended in PBS and quantified different sets either using BD FACS Aria ^TM^ (BD Biosciences) or BD LSR II (BD Biosciences) flow cytometer and data were analysed using BD FACS DIVA ^TM^ (BD Bioscienc es) flow cytometer.

### Superresolution SIM images

All SIM (structured illumination microscopy) images were acquired using on DeltaVision OMX SR microscope from GE Healthcare using a 60X 1.42 NA objective and pco.edge sCMOS detector. The cells were fixed by 4% PFA by incubation for 15 min and fixed cells were imaged. The widefield images were deconvolved using the built-in algorithm.

### Image analysis and statistics

Image analysis was done using ImageJ software (open source by NIH, USA). Percentage of co-localization of oAβ (magenta) with lysosomes (green) was performed by calculating the proportion of the magenta fluorescence pixels compared to the co-localized pixels from the background subtracted images using the Coloc-2 plugin. The number of TNTs were distinguished from neurites by comprising a 3D-volume view of cells from confocal stacks using the volume view plugin in Image J software. Cells with blebs / lamellipodia and TNTs were counted from images and normalized to the total number of cells and represented in percentage. oAβ induced endocytosis was quantified by measuring the integrated intensities of internalized TMA-DPH from the luminal part of each cell by drawing ROI (region of interest) over sequences of time-laps confocal image stacks and comparing it with the same quantification of control cells. The fusion of lysosomal membrane to reseal the damaged membrane was detected as the appearance of LAMP1 (green) on the outer leaflet of the PM. The outer leaflet LAMP1 was quantified measuring the integrated intensities from drawn ROI and the proportionate percentage calculated comparing the total LAMP1 per cells. One-way ANOVA tests were performed to validate statistical significance in all experiments.

## Results

### oAβ induced cellular stress instigates formation of TNTs and cell-to-cell pathology propagation

Cell-to-cell propagation of oAβ between connected cells has been shown in earlier studies ^4, 5, 8, 14^. In our previous work ^4^, we have shown asymmetric spreading of oAβ along the dispersion of neurons path, when microinjected in a single neuron of a primary hippocampal culture ^4^. Additionally, we showed transfer of oAβ-TMR from connected donor to acceptor cells using a 3D donor-acceptor co-culture model with differentiated SH-SY5Y cells. However, the clear mechanism behind the cell-to-cell propagation remains to be revealed. Differentiated cells both in 2D and 3D culture form neurites and express several neuronal markers and characteristics of mature neurons, where 3D differentiated cells express tau subtypes better comparable to mature human neurons than 2D differentiated ^4, 22^. Here, we have observed that differentiated cells in 2D culture upon treatment with oAβ-TMR, morphologically exhibit lamellipodia, cell membrane expansion or blebs (yellow arrows) as well as tunneling nanotube-like (TNT-like) long thin conduits (white arrows) between neighbouring cells (Fig. 1B, Supplementary Movie 1). We did not observe significant numbers of TNT-like structures and membrane expansions/lamellipodia in the differentiated control cells (Fig. 1A). We have quantified the number of cells with blebs/lamellipodia and counted the number of TNT-like structures from bright field images with respect to the total number of cells from each of the image frames and presented as a percentage. The results show a concentration-dependent increase in oAβ-TMR treated cells (200-500 nM) compared to the control cells (Fig. 1C, D). We have detected the direct transfer of oAβ-TMR between the cells prominently (indicated by blue arrows) via these TNT-like conduits (Fig. 1E-F, Supplementary Movie 3-4) by time-lapse imaging. Transfer of organelles and oAβ-TMR (blue arrows) via TNT-like conduits extended from lamellipodia between cells (Yellow arrows), were observed in the cells incubated for 3 h with 100, 250 and 500 nM of oAβ-TMR (Supplementary Movies 2-4).

**Figure 1:**
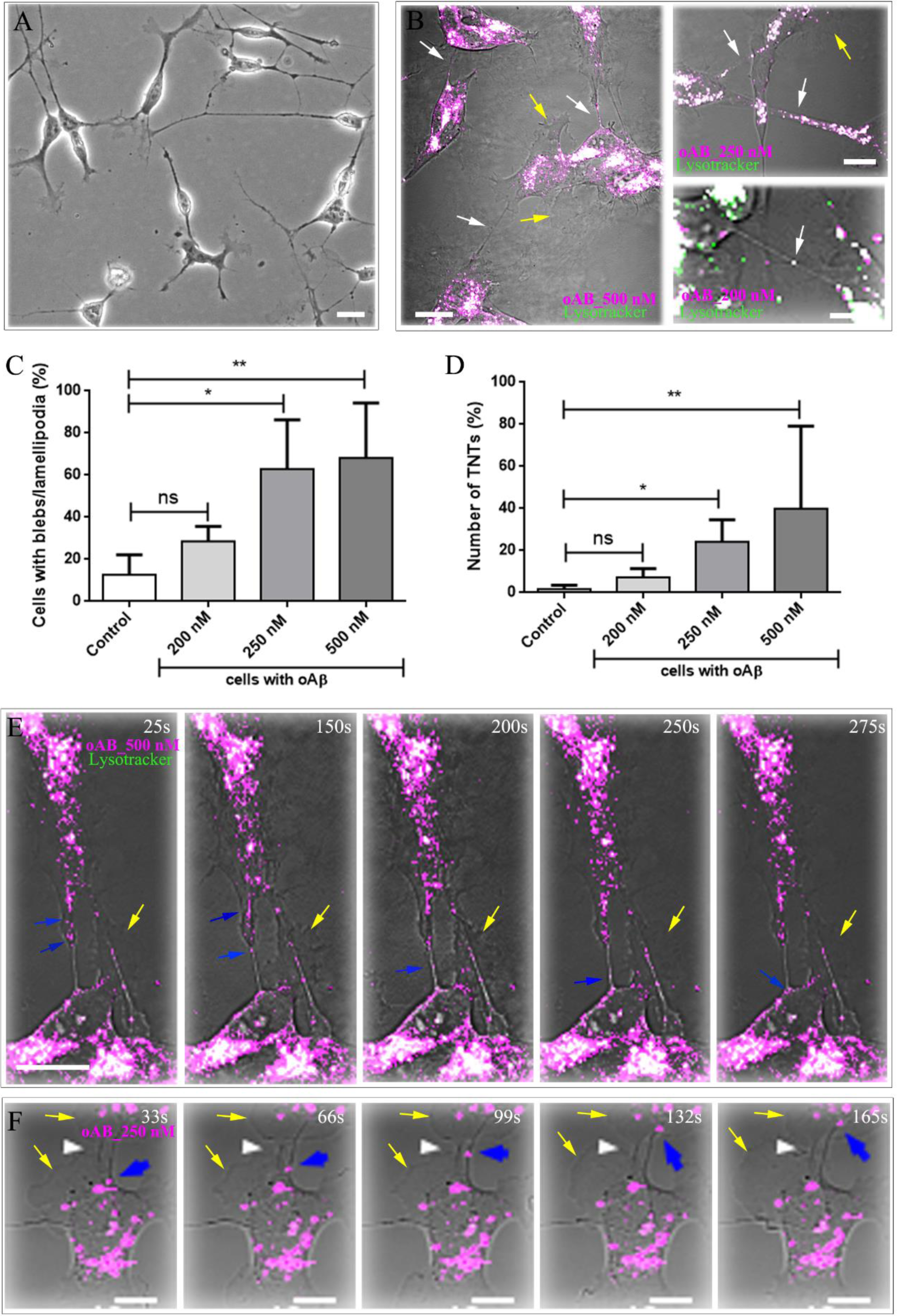
oAβ induced formation of TNT-like conduits in differentiated SH-SY5Y neuronal cells. A-B) TNT-like conduits (indicated by white arrows) between neighbouring cells were detected in differentiated SH-SY5Y cells incubated with 500, 250 and 200 nM of oAβ-TMR (magenta) for 3 h, washed and stained with 50 nM of lysotracker (green) before the capture of the image. The cells with long TNT-like conduits form noticeable blebs and cell membrane expansion (indicated by yellow arrows), in contrast to control cells. A) Differentiated control cells showed neurite like structures rather than TNT-like conduits. Note that the control cells are also devoid of blebs and cell membrane expansion. C) Percentages of cells with blebs / lamellipodia were quantified from the images taken with increasing concentrations of oAβ-TMR (200-500 nM) and compared with the control cells. D) TNT-like conduits were counted and plotted in percentage with respect to the number of cells from each of the image frames. Quantifications were done from > 60 cells in each set. n > 3. Plots are mean ± SD. One-way ANOVA tests were performed to validate statistical significance. E) Differentiated SH-SY5Y cells were incubated with 500 nM of oAβ-TMR (magenta) for 3 h, washed and labelled with 50 nM of lysotracker (green) and a sequence of images at different time points were taken. The cells form TNT-like conduits and oAβ-TMR travels from one cell- to-another as organelle like puncta structures through the conduits (blue arrows). The cells form noticeable blebs (yellow arrows). F) The cells treated with 250 nM oAβ-TMR (magenta) for 3 h form lamellipodia (yellow arrows), show direct transfer of oAβ-TMR from one cell-to- another as organelle like puncta structures (blue arrows) through the TNT-like conduits (white arrows). Scale bars are 10 μm.

In order to show that the studied TNT-like structures are indeed oAβ induced TNTs and not an artefact of differentiating reagents, we performed the experiments in partially differentiated (treated with retinoic acid for 7 days) and undifferentiated SH-SY5Y cells in parallel and confirmed that TNTs from 3D volumetric images obtained from confocal z-stacks. We have observed thin TNT-like structures extended from expanded lamellipodia-like membrane protrusions in partially differentiated cells internalized with oAβ-TMR after incubation with 250 nM of oligomers for 3 h (Fig. 2A-B), in contrast to the control cells (Fig. 2C). Moreover, the partially differentiated SH-SY5Y cells formed networks of TNT-like conduits between neighbouring cells (yellow arrow, Fig. 2B). The cells make networks between 3 neighbouring cells via TNT-like conduits (yellow arrows), and those were also extended from expanded lamellipodia (black arrows, Fig. 2A). We have observed the transfer of oAβ-TMR colocalized with Lysotracker labelled lysosomes (yellow arrows) from one cell to another via the TNT-like conduits extended from expanded lamellipodia (black arrows; Fig. 2D). To confirm that the cell-to-cell conduits visible in bright field images are TNTs, we have composed 3D volume view from confocal z-stacks of phalloidin stained f-actin structures between neighbouring cells (yellow arrows, Fig. 2E-F). In the 3D volume view, TNTs were distinguished from neurites by the characteristics of their capacity to stay hanging without touching the substratum even after fixing the cells (Fig. 2F). The lengths of the TNTs were between 0.2-16 μm. Diameters of TNTs were measured from confocal z-stacks images of phalloidin stained cells and the values of majority (> 90%) of TNTs are within 240-960 nm measured at xy-plan (between 1-4 pixels, pixel size is 240 nm). However, around 10 % of TNTs are thicker and their diameters are in the range of 1.23 ± 0.27 μm. Finally, we have quantified the number of TNTs and neurites per cell in partially differentiated cells treated with 1 μM of oAβ-TMR over time (1-3 h), and compared to the control cells (Fig. 2G-H). The results show increasing numbers of TNTs over the time (1-3 h) of oAβ (1 μM) treatment (Fig. 2G). At the same time, we observed a significant decrease in the number of neurites per cells after 3 h of oAβ (1 μM) treatment (Fig. 2H). Similarly, oAβ showed toxicity to the neurites of differentiated cells, which caused significantly reduce number of neurites per cell detected in a concentration and time dependent manner (Supplementary Fig. 1A-D).

**Figure 2:**
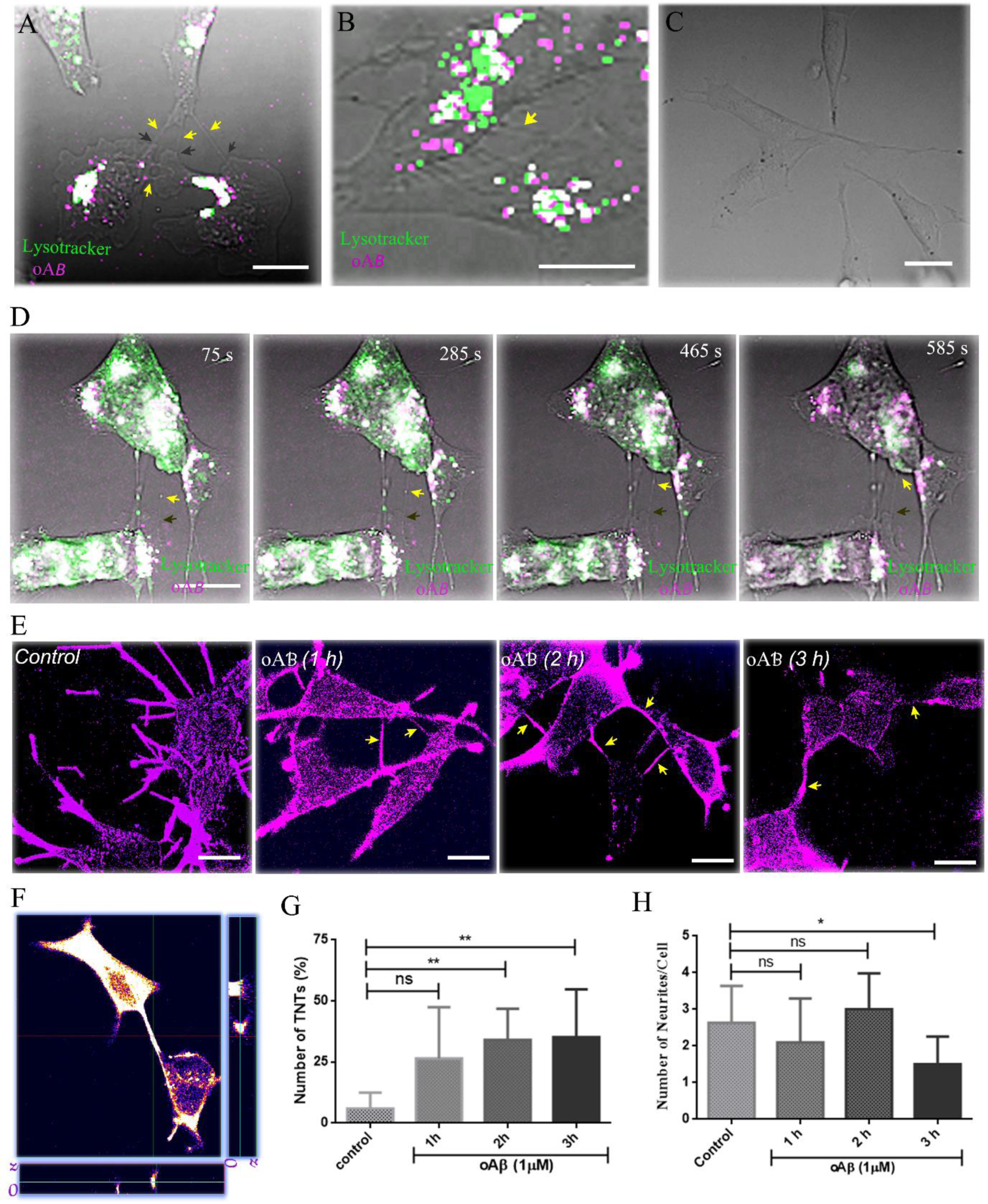
oAβ induced formation of TNTs in partially differentiated SH-SY5Y neuronal cells. A-B) Partially differentiated SH-SY5Y cells were incubated with 250 nM of oAβ-TMR (magenta) for 3 h, washed and labelled with 200 nM of lysotracker (green). A) oAβ- TMR internalized cell makes network between 3 neighbouring cells via formation of TNT-like connections (yellow arrows), extended from lamellipodia-like membrane protrusions (black arrows). B) TNT-like conduits form networks between two neighbouring cells (yellow arrow). C) Partially differentiated control cells devoid of blebs / lamellipodia. D) Partially differentiated SH-SY5Y cells were incubated with 250 nM of oAβ-TMR (magenta) for 3 h, washed and labelled with 200 nM of lysotracker (green). The cells form thin TNT-like conduits (yellow arrows) extended from expanded lamellipodia-like membrane protrusions (black arrows). oAβ-TMR co-localized with lysosomes that moves from one cell to another via connected conduits (yellow arrows). E-F) 3D volume views were composed from confocal z- stacks of phalloidin stained f-actin structures between neighbouring cells to validate whether the cell-to-cell conduits which were visible in bright field images are TNTs or not. G-H) Then, TNTs were distinguished from neurites from the characteristics of their capacity to stay hanging without touching the substratum even after fixing the cells. Quantifications of G) TNTs and H) neurites were done in the cells treated with 1 μM of oAβ for 1-3 h, compared to the control cells, from > 60 cells in each set. Plots are mean ± SD. One-way ANOVA tests were performed to validate statistical significance. Scale bars are 10 μm.

### oAβ induced TNT formation precedes enhanced membrane activities and massive endocytosis

We have designed the experiments using undifferentiated SH-SY5Y cells to ensure that it is the toxicity of oAβ that causes induction of TNT-like structures, and not the differentiating reagents. Cells were co-transfected with lifeact-EGFP and CAAX-mCherry, to stain actin and peripheral membrane proteins. We found formation of numerous co-stained TNT-like structures (yellow arrows) after 1h of oAβ (1 μM) treatment, compared to the control cells (Fig. 3A-B). Further, the TNT-like structures were confirmed as a continuous extension of the PM and actin protrusions between two cells from 3D volume view from confocal z- stacks (Fig. 5B). Preceding the formation of oAβ induced TNTs, we have observed enhanced membrane activities along with the formation of membrane ruffles, filopodium, blebs and massive endocytosis compared to the control cells (Fig. 3C; Supplementary Movies 5-6). Enhanced membrane activities and endocytosis were detected immediately at the addition of oAβ (1 μM). The oAβ-induced endocytosis was quantified by measuring the internalization of TMA-DPH into the luminal part of the cells, compared to the control cells (Fig. 3D). TMA- DPH is membrane impermeable and able to enter the control cells and to the cells treated with oAβ (1 μM, for 15 min) only via endocytosis. On the other hand, the membrane impermeable dye TMA-DPH is able to enter the cells treated with oAβ (1 μM) for 1 h, and the dye stains the whole cells (oAβ treated for 1 h) immediately at the addition (Fig. 3E). The result suggests that oAβ induced changes in the membrane fluidity.

**Figure 3:**
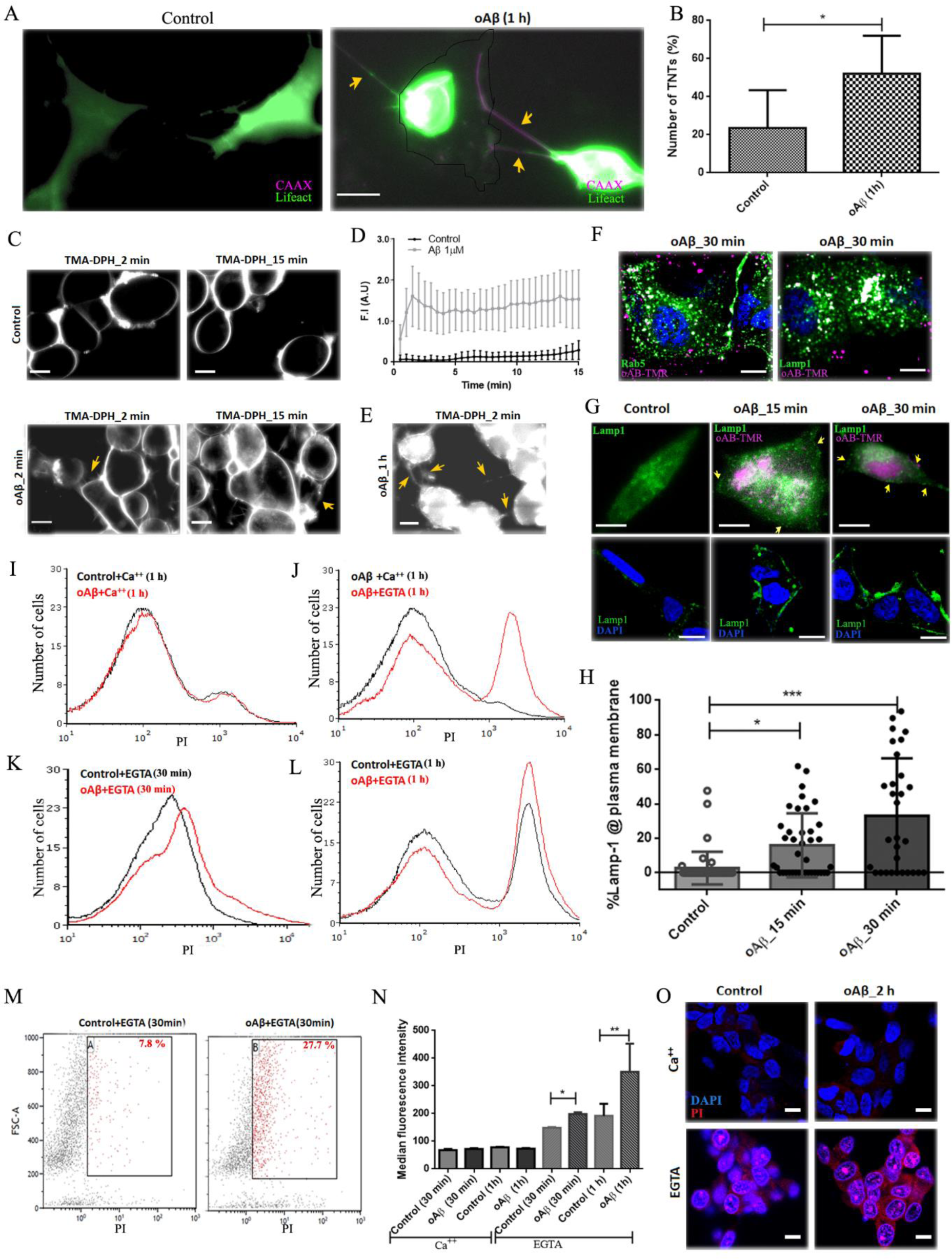
oAβ induced PM damage and repair via coupled lysosomal-exocytosis and endocytosis. A) TNT-like conduits (yellow arrows) were quantified in undifferentiated SHSY5Y cells co-transfected with lifeact-EGFP (actin) and CAAX-mCherry (peripheral membrane-protein). B) Quantification of TNT-like conduits from number of cells > 60 for each set. C) Massive membrane activities and endocytosis were observed in the oAβ (1 μM) treated cells together with the membrane dye TMA-DPH (0.5 μM), compared to control cells. D) Endocytosis of PM labelled with TMA-DPH was quantified by measuring luminal part intensities (plotted mean ± SD, quantified > 20 cells from each set, n=3). E) In oAβ (1h with 1 μM) treated cells, penetration of the membrane impermeable dye TMA-DPH on addition, and the cells show TNT-like conduits (yellow arrows). F) Cells incubated with extracellularly applied oAβ-TMR (1 μM) internalize (magenta) efficiently to early-endosomes (Rab 5; green) and late-endosomes/lysosomes (Lamp1, green). G) Translocation of Lamp1-mGFP to cell surface (Upper panels), n=3. H) Lamp1 surface staining (Lower panel) within 15-30 min of exposure of oAβ (1 μM) was quantified from intensity (plotted mean ± SD) measurements from defined ROI using ImageJ. (n=3, each dot represents the number of cells). I-J) oAβ (1 μM) induced membrane damage detected as the uptake of the membrane-impermeable dye PI in presence and absence of Ca^2+^, were quantified by flowcytometry. Represented histograms I) In presence and K-L) absence of Ca^2+^. M) Representative dot-plot in absence of Ca^2+^. N) Penetration of PI presented as a change of median fluorescence in control and oAβ (1 μM, 30 min and 1 h) treated cells in presence and absence of Ca^2+^ (n = 6). O) Representative confocal images of PI uptake after 2 h of oAβ treatment compared to control. Plotted mean ± SD. One- way ANOVA were performed to validate statistical significance. Scale bars are 10 μm.

To understand if the oAβ induced TNT formation is specific to neuronal cells or if it is a basic cellular process, we incubated HEK-293 cells with oAβ (1 μM) for 1 h and observed formation of long TNTs hanging between distant neighbouring cells (Supplementary Fig. 2C). The number of TNT like conduits were significantly lower in control cells (Supplementary Fig. 2A-B). Similar to SH-SY5Y cells, the membrane of control HEK-293 cells found to be labelled by TMA-DPH only at the periphery (Supplementary Fig. 2D). However, in the oAβ (1 μM for 1 h) treated HEK-293 cells, TMA-DPH labelled whole cells by entering the cells immediately on addition. Additionally, these cells also showed TNT-like protrusions (Supplementary Fig. 2E). Results suggest oAβ induced change in membrane fluidity and increased number of TNT-like structures.

### oAβ1-42 induces membrane damage and lysosomal exocytosis

oAβ induces TNT formation in association with the substantial enhancement of membrane activities and massive endocytosis, similarly as evident in Ca^2+^ dependent repair of injured PM by lysosomal exocytosis ^28^. We have observed that the oAβ-TMR (magenta) was efficiently internalized into early endosomes (Rab5 positive organelles), followed by entry into multivesicular bodies (MVB) or lysosomes (Lamp1 positive organelles) (Fig. 3F). The spontaneous internalizations of extracellularly applied oAβ-TMR (250 nM to 1 μM, incubated for 15 min to 1 h) into undifferentiated cells were quantified and results showed that 70 ± 9 % of oAβ-TMR (250 nM) ended up in Lamp1 positive organelles after 15 min of incubation (Supplementary Fig. 3A-B). Images in Fig. 3F, present the colocalization of oAβ-TMR (1 μM) after 30 min of incubation. This inspired us to determine if oAβ caused PM damage. The Gold standard to detect PM repair by lysosomal exocytosis and formation of a patch over the damaged membrane is to detect the luminal part of the lysosomal membrane protein LAMP-1 exposed on the outer leaflet of the plasma membrane. Accordingly, we have observed transiently transfected Lamp1-mGFP distributed in large extent in the periphery of the PM within 15 to 30 min of oAβ (1 μM) treatment, compared to the control cells (Fig. 3G upper panel). We have quantified oAβ (1 μM) induced Lamp1 on the PM, in undifferentiated SH-SY5Y cells within 15-30 min of exposure by surface staining of the cells (Fig. 3G lower panel & Fig. 3H). To verify that the process is calcium dependent, we analysed the influx of the membrane-impermeant dye propidium iodide (PI) in presence of 5 mM EGTA in PBS. The rationale behind this experiment is that if lysosomal exocytosis-dependent PM repair is occurring, then chelation of Ca^2+^ will prevent the repair and PI will be detected intracellularly. As seen in Fig. 3I-M, significant enhancement of PI staining in absence of calcium was detected 30 min, 1h and 2 h after exposure to oAβ as quantified by flow cytometry (Fig. 3K-M) and confocal imaging (Fig.3O). In the presence of Ca^2+^ increased PI staining was not detected with oAβ treatments (Fig. 3I-J). The quantification of internalized PI was compared by plotting median fluorescence intensity, as fluorescence histogram profiles are bimodal in nature (Fig.3N). The presented dot plot of PI internalization after 30 min of Ca^2+^ chelation in the presence and absence of oAβ (1 μM) (Fig. 3M). The results show an increasing amount of internalization of PI with time even in the control cells, however the differences between control and oAβ (1 μM) treated cells are significant at both 30 min and 1 h time point, and even at 2 h time point differences are detectable. The results suggest that Ca^2+^ dependent membrane repair is a continuous process even in the control cells, and oAβ induced membrane damage accelerates the process. Thus, we conclude that the addition of oAβ causes damage to the PM and as a result of rapid membrane repair process occurs, by enhancing the process of lysosomal exocytosis and fusion of lysosomal membrane with the PM. Subsequent to the membrane repair process, re-establishment of the PM occurs by removing membrane parts through endocytosis and as a consequence, oAβ is internalized into MVB/lysosomes (Lamp1 positive organelles) (Fig. 3F and Supplementary Fig. 3A-B).

### Involvement of the actin cytoskeleton remodeling kinase PAK1 engaged endocytosis in the internalization of oAβ1-42

We next wanted to determine the exact mechanism of the massive internalization of oAβ. To identify the mechanisms of internalization of oAβ, partially differentiated SH-SY5Y cells were pre-treated with different inhibitors against the uptake of Aβ as suggested in the literature. We have used partially differentiated cells, because N-methyl- D-aspartate (NMDA) receptors, a-amino-3-hydroxy-5-methyl-4-isoxazolepropionic acid (AMPA) receptors and nicotinic acetylcholine receptors (nAChR) are known to be present on RA treated partially differentiated SH-SY5Y cells ^29^ and have previously been shown to mediate the internalization of Aβ ^30–32^. To inhibit the function of these receptors, antagonists against NMDA receptors (AP-5), AMPA receptors (GYKI 52466) and nAChR (α- Bungaratoxin) were used and oAβ-TMR uptake was quantified by flow cytometry. However, no change in the uptake of oAβ was found (Fig. 4A). To determine the mechanism of Aβ-uptake several inhibitors that affect different variants of endocytosis and macropinocytosis were tested. After pre-treatment with the inhibitors, the cells were treated with oAβ-TMR and quantified by flow cytometry (Fig. 4B-C). The percentage of oAβ-positive cells was normalized to the mean percentage of oAβ-uptake and the fold-change upon inhibitor treatment was calculated. MDA and PAO, which are inhibitors of clathrin- mediated endocytosis, ^33^ and DPA, which functionally inhibits acid sphingomyelinases ^34^ could not prevent the uptake. Similarly, the dynamin inhibitor dynasore ^35^, and an amiloride analogue, EIPA that acts as a specific inhibitor of macropinocytosis through inhibition of Na^+^/H^+^ exchangers ^36^ had no effect. However, we found significantly reduced oAβ-uptake when using bafilomycin A1 (Baf) and NH4Cl, which both affect the lysosomal acidification and consequently fusing between vesicles ^37^ and the PAK1 inhibitor IPA-3 that inhibits actin regulated endocytosis similar as macropinocytosis by modulating PAK1 dependent actin polymerization ^38, 39^. PAK1 has an autoregulatory domain which is targeted by the inhibitor IPA-3, although the exact downstream inhibitory signalling pathway of IPA-3 is not known yet ^38^. To establish if the oAβ internalization in undifferentiated SH-SY5Y cells occurs similarly via PAK1 dependent endocytosis, we also quantified the internalization in undifferentiated cells and the quantification showed similar results, Fig. 4D shows the representative confocal images. Involvement of PAK1 has been reported mostly in membrane tension and cholesterol sensitive clathrin-independent endocytosis (CIE) and in membrane modulating actin dependent endocytosis such as, macropinocytosis, IL2Rβ endocytosis, CLIC/GEEC (clathrin-independent carrier/ glycosylphosphatidylinositol-anchored protein enriched compartment) pathway ^39, 40^. It has been shown that PAK1 specifically regulates macropinocytosis by following the uptake of 70k Da dextran, a specific marker of fluid phase macropinocytosis ^41^. Therefore, we have followed the fate of internalization of Dextran-70 kDa with oAβ pulses. However, after 1 h of oAβ-TMR (1 μM) treatment only small amounts of internalized dextran-FITC were detectable and there was no significant colocalization with internalized oAβ-TMR (Fig. 4E). Similarly, internalizations of oAβ were observed in the GFP-GPI (GFP- glycosylphosphatidylinositol) transiently transfected SH-SY5Y cells, and the internalized oligomers were not colocalized with GFP-GPI in the cells treated for 1 h with oAβ-TMR (1 μM) (Fig. 4E) and oligomers did not enhance the GFP-GPI internalization observed after 15 - 30 min of oAβ (0.5 nM to 1 μM) treatment. The results indicate that oAβ follows PAK1 dependent CIE machineries, which is distinct from macropinocytosis or the CLIC/GEEC pathway. The same observations were also reported recently ^39^ although the complete mechanism is not fully understood.

**Figure 4:**
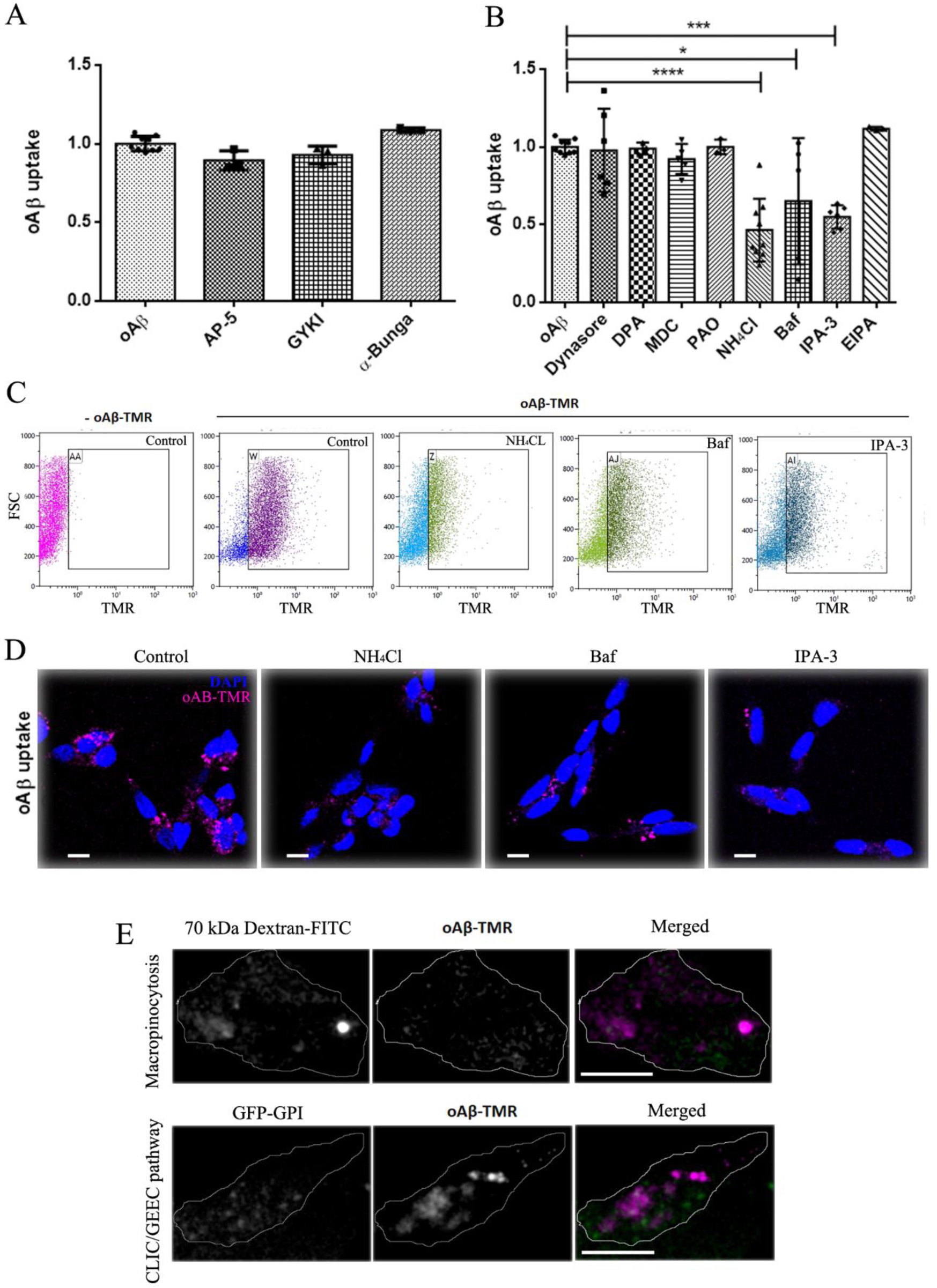
Mechanisms of rapid internalization of oAβ-TMR were identified by pre-treating the SH-SY5Y cells with different endocytosis and receptor inhibitors. The percentages of oAβ-TMR containing cells in partially differentiated cells were analysed by flow cytometry. A) Cells were pretreated with the receptor inhibitors α-Bungarotoxin, AP-5, GYKI. n=1, triplicates. B) Cells were pretreated with endocytosis inhibitors Dynasore, DPA, MDC, PAO, NH4Cl, Baf, IPA-3 and EIPA, followed by exposure to 0.5 μm oAβ-TMR for 1h. n= 3, > minimum of duplicates in each set. The percentage values were normalized to the respective mean of oAβ. C) Representative dot-plots with forward scatter vs fluorescence intensities (TMR). D) Representative confocal images of undifferentiated SH-SY5Y cells treated with endocytosis inhibitors and oAβ-TMR. E) Representative images are showing that internalized oAβ-TMR (magenta; 1 μM incubated for 1 h) was not colocalized with 70 kDa Dextran-FITC (green) and GFP-GPI (green). One-way ANOVA tests were performed to validate statistical significance. Scale bar = 10 μm.

**Figure 5:**
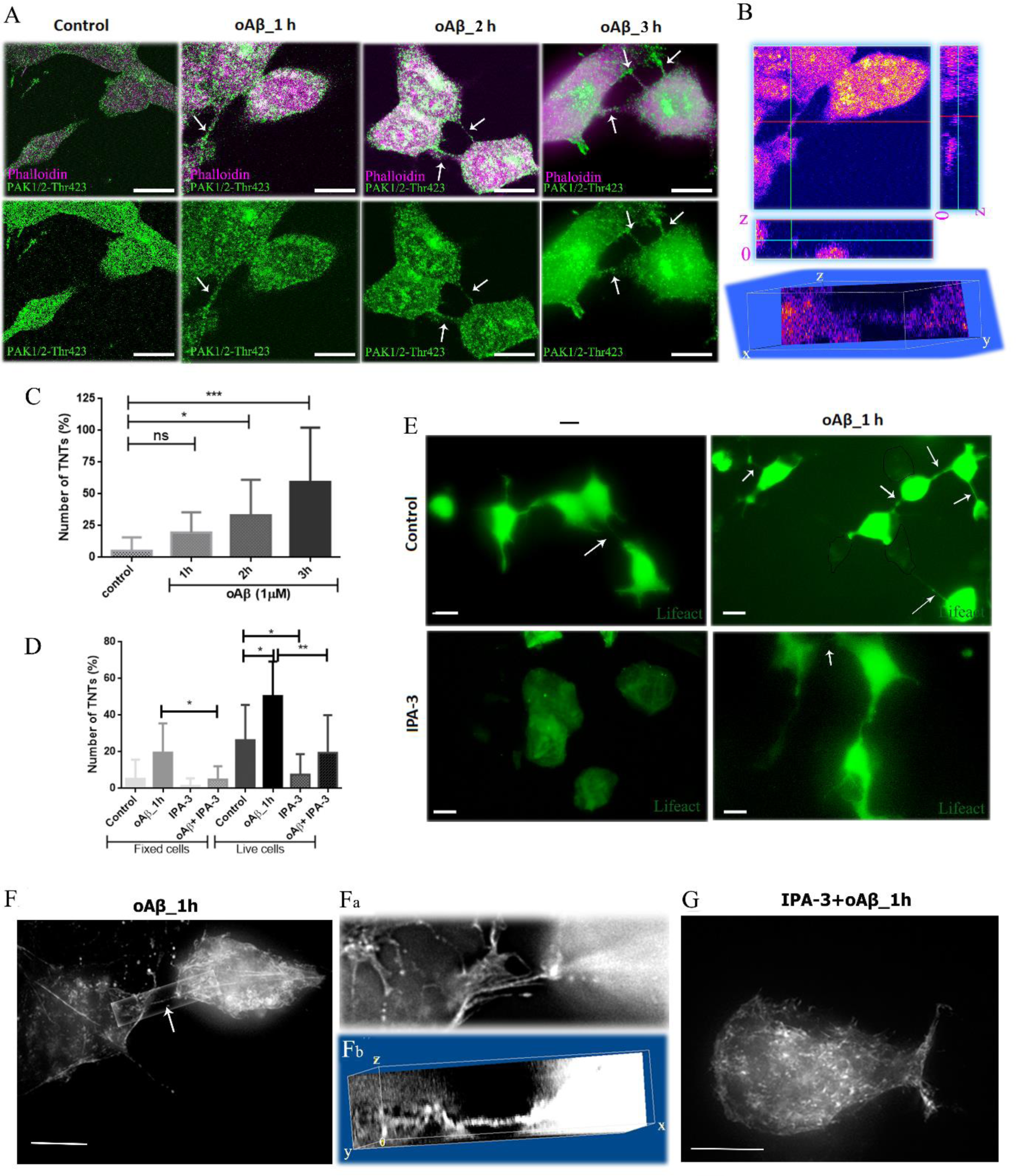
Inhibition of rapid internalization of oAβ by the PAK 1 inhibitor IPA- 3 can also inhibit formation of TNTs. A) Represented oAβ induced TNTs from confocal z- stack images in the undifferentiated SH-SY5Y cells co-stained by phalloidin (magenta) and anti PAK1/2-Thr423 (green). B) TNTs were quantified creating 3D volume view from z-stacks confocal images. C) Quantification of TNTs in the cells treated with 1 μM of oAβ for 1-3 h, compared to the control cells. E) Fluorescence microscopy images of control cells, oAβ (1 μM) treated, IPA-3 treated and cells treated with IPA-3 and then oAβ were captured using undifferentiated SH-SY5Y cells transfected with lifeact EGFP (green) (white arrows). D) Number of TNTs were counted with respect to the number of cells to quantify the inhibition by IPA-3 and compared the TNT numbers both in live cells and fixed cells. Number of cells > 50 for each set. Plotted mean ± SD. One-way ANOVA tests were performed to validate statistical significance. F) Superresolution images of lifeact stained undifferentiated SH-SY5Y cells by DeltaVision™ OMX SR microscopy. SH-SY5Y cells were transfected with Plenti- lifeact EGFP and treated with oAβ (1µM) and IPA-3 (20µM). F) In oAβ (1 h) treated cells, TNTs appear to spread out as extensions from the long f-actin of the cytoskeleton and to connect the two neighbouring cells. Fa) Zoomed image of a proper z-stack illustrated clearly the f-actin labelled TNT. Fb) 3D volume view of z-stacks of SIM image shows that, in contrast to neurons, TNTs do not touch the substratum, but rather stay “hanging” between the two connected cells even after the fixation of the cells. G) In the IPA-3+oAβ treated cells, disruptions of long stretched f-actin of cytoskeleton and inhibition of TNTs are evident. 3D reconstructions and analysis were done using ImageJ. Scale bars are 10 μm.

### Active PAK1 in the oAβ induced formation of TNTs

ActivePAK1 regulates cortical actin polymerization, directional movements and also polarized lamellipodia at the leading edge ^42^. Actin depolymerizing drugs such as latrunculin and cytochalasin impair TNTs formation ^17^. Therefore, the next step was to observe the role of PAK1 in the oAβ induced formation of TNTs and if inhibition of oAβ uptake by IPA-3 also prevented the formation of TNTs. Thus, oAβ induced formation of TNTs in undifferentiated SH-SY5Y cells was quantified by creating 3D volume view from z-stacks images (Fig. 5B) of phalloidin and anti PAK1/2-Thr423 stained conduits between neighbouring cells at fixed condition (Fig. 5A). Number of TNTs was quantified in cells treated with 1 μM of oAβ for 1-3 h, compared to the control cells (Fig. 5C) and results show a significant time dependent increase in the number of TNTs after oAβ treatment. Paraformaldehyde fixation might break or damage the thin TNTs. Thus, TNTs were also quantified by staining f-actin in live cells by transiently transfecting with the lifeact-EGFP plasmid (Fig. 5E). Transfection did not affect the ability of cell to form TNT-like structures and the increased number of TNT-like structures in oAβ treated cells as compared to the controls maintained. Cells that were treated with IPA-3 showed a decrease in number of TNTs (Fig. 5D-E), images were quantified from live cells as well as from fixed condition (Fig. 5D). The quantification of TNTs in the live cells was higher but the pattern is similar to the fixed cells, which were quantified from phalloidin stained TNTs images by analysing in 3D volume view from the confocal Z-stacks (Fig. 5E). Morphologically, IPA-3 treated cells were rounder than the controls and oAβ treated cells, however MTT assay showed no significant change in the cell viability (Supplementary Fig. 4A).

### oAβ induced activation of phospho-PAK1 modulates f-actin and formation of TNTs

Distinct differences in f-actin structures between oAβ treated and the oAβ + IPA-3 treated cells were detected by observing the lifeact EGFP plasmid labelled f-actin structures at a resolution of < 130 nm using Superresolution SIM (Structured Illumination Microscopy) images. TNTs were observed to be a continuous extension of long cortical f-actin in oAβ treated cells (Fig. 5F and Fa). 3D volume view of the xyz-plane demonstrated that membrane actin protrusion did not grow on the surface like neurites. Rather, they were ‘hanging’ between two neighbouring cells, which is one of the characteristics distinguishing features of TNTs (Fig. 5Fb). In the Fig.5G, the disruptions of long actin fibres in IPA-3 pre-treated cells reveal PAK1 play an important role in f-actin modulation and thereby TNT formation. Experiments were done with undifferentiated sparsely seeded SH-SY5Y cells to quantify actin stained TNTs like structures between relatively distant cells (Supplementary Fig. 4B).

Images of immunocytochemical staining using phospho-PAK1 (Thr423)/PAK2 (Thr402) antibody demonstrated increased levels of activated phospho-PAK1 with time (1-3 h) of oAβ (1μM) treatment (Fig. 6A-B). The confocal images were represented as z-projected stacks and the intensities were compared from z-projected images, taken at the same settings of laser power and exposure. Co-staining of activated phospho-PAK1 with f-actin on TNT structures were confirmed from the 3D-volume view analysis (Fig. 5A-B & 6A). Upregulation of activated phospho-PAK1 by oAβ (1μM) treatment was quantified further from epi- fluorescence images, where images were taken from more than 80 cells from each set using 20X magnification objective and at the same exposure and settings (Fig. 6C). The results show a higher level of activated phospho-PAK1 in oAβ (1μM) after 1 h treatment of cells in comparison to the control and IPA-3 pretreated cells (Fig. 6D). Further quantifications were done by flow cytometry using immunocytochemical stained cells. The results obtained from the histograms of flow cytometeric data showed higher expression of phospho-PAK1 and phalloidin bound actin in oAβ treated cells as compared to control, IPA-3 and IPA-3 + oAβ treated cells (Fig. 6F-G). Quantification of TNT-like cell-to-cell phalloidin stained actin extensions with respect to the level of activated phospho-PAK1 showed a positive correlation when quantified observing a larger number of cells (Fig. 6E).

**Figure 6:**
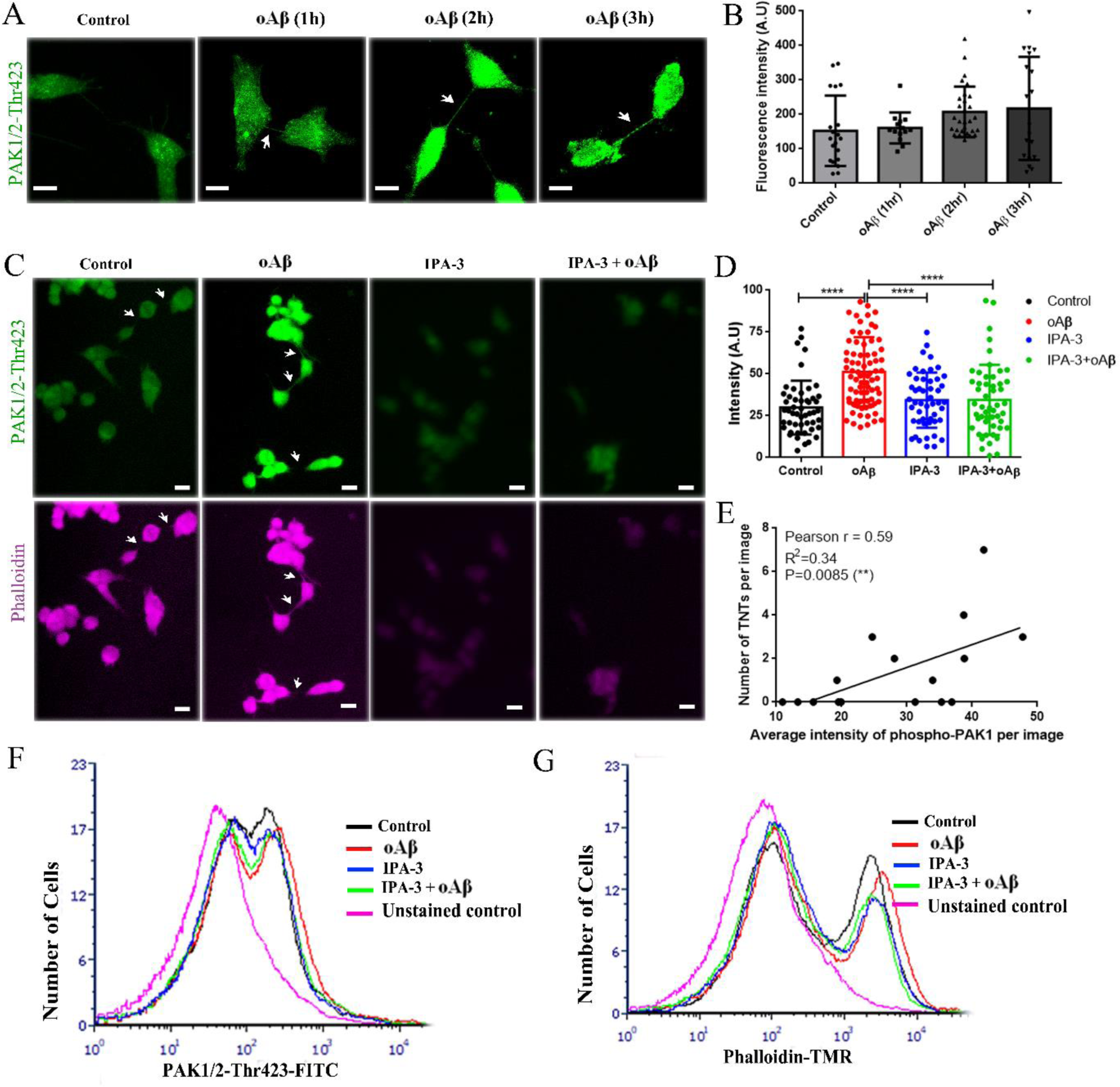
oAβ induced activated phospho-PAK1 modulates f-actin extensions and IPA-3 disrupts the long-stretched f-actin. A) Confocal images of immuno-stained cells treated with oAβ (1 μM) over the time show localization of phospho-PAK1 (Thr423)/PAK2 (Thr402) antibody (green) on TNTs (indicated by white arrows). B) Intensity analysis of z-projected confocal images show increase level of activated phospho-PAK1. C) Images of immuno-stained cells using phospho-PAK1 (Thr423)/PAK2 (Thr402) antibody (green) and actin-binding phalloidin-TMR (magenta) were taken by 20X 0.4 NA objective. D) Level of active phospho-PAK1 were quantified measuring the intensities from each cell using image-j ROI manager after background subtractions (number of cells >80 in each set) with oAβ, IPA- 3 and IPA-3+oAβ treated cells. E) Number of TNT like cell-to-cell connections (indicated by white arrows) show a positive correlation with the intensities of activated phospho-PAK. F-G) Level of active phospho-PAK1 and phalloidin labelled actin were quantified by flowcytometry in the cells treated with oAβ, IPA-3 and IPA-3+oAβ in compared to the control. One-way ANOVA tests were performed to validate statistical significance. Scale bars = 10 μm.

## Discussion

Prion-like cell-to-cell propagation is a common characteristic of neurodegenerative diseases. Several reports have consistently reported direct cell-to-cell propagation of neurodegenerative protein aggregates and their implications in the gradual pathology progression ^2, 3^. Several studies have suggested exosomes as a means of cell-to-cell transfer of Aβ ^13, 14^. On the other hand, studies using the method of transwell assays have also shown the efficient cell-to-cell transfer of prion proteins despite the blocking of exosome transfer ^9, 43^. Moreover, an increasing number of reports show that cell-to-cell transfer of neurodegenerative proteins, such as PrP^Sc^, α-synuclein, tau, polyQ aggregates and Aβ, via TNTs instigate new avenues ^8–11^. oAβ induced formation of TNTs in primary neurons and astrocytes has already been reported ^8^. Here we have focused more on possible mechanism of oAβ induced formation of TNTs in differentially differentiated SH-SY5Y neuronal cells. Due to its cancerous origin, the SH-SY5Y cell line shows a number of genetic aberrations and different differentiation protocols generate variations in neuronal properties, but most genes and pathways dysregulated in AD pathogenesis stay intact even in undifferentiated SHSY5Y cells. Moreover, its differentiation to cholinergic neurons like properties in retinoic acid treated SHSY5Y cells a widely accepted model system in AD research ^22^.

Increasingly, clinical studies and animal models indicate that soluble oAβ is the disease initiator and driver, rather than large extracellular depositions. Accumulation of amyloidogenic proteins in lysosomes, abnormal lysosomal morphology and lysosomal membrane permeabilization are major hallmark of neurodegenerative diseases. Notably, lysosomal stress as well as damage due to accumulation of non-degradable amyloidogenic aggregates could induce the formation of TNTs ^12^. Stress signals from lysosomes dysregulate various cellular processes and mediate increased oxidative stress ^12, 44–46^. Several studies have indicated that ROS (reactive oxygen species) induced cellular stress enhances TNT formation ^8, 17^.

In contrast, the study ^47^, demonstrated that the impaired processing of Aβ due to lysosomal enzymatic inefficiency can enhance exocytosis. Additionally, a related protein of the exocyst complex M-sec, involved in exosome fusion and membrane expansion, regulates formation of TNTs ^21^. PM recruitment of Ral-GTPase and filamin, both actin remodeling proteins, also indicate positive regulating effects in TNT formation ^21^. The study presented here also demonstrates membrane expansion in the form of blebs, filopodium-like structures and extension of TNTs from expanded lamellipodia. Previous reports have indicated that synthetic oAβ makes ion-permeable pores in synthetic membranes ^48, 49^. Recently, it was also shown that oAβ can induce a membrane repair response similar to that induced by exposure to the bacterial pore-forming toxin produced by *B. thuringensis* ^50^. Consequently, enhanced internalization of Aβ occurs via endocytosis, which is independent of receptor interactions. Involvement of the clathrin- and dynamin-independent endocytosis is also relevant in maintaining cellular homeostasis by regulating membrane stress and cell surface expansion ^39, 51^. A recent study reported that Aβ follows membrane tension sensitive and Rho GTPase family regulated actin- dependent CIE ^39^. Furthermore, Aβ uptake was earlier shown to be inhibited by nocodazole and cytochalasin-D, the inhibitors of tubulin depolymerization and actin polymerization, by preventing fluid phase-endocytosis in microglia and astrocytoma cells ^52^. The results of this study show that the internalization of oAβ through massive endocytosis is clathrin- independent, but rather follows PAK1-dependent membrane actin modulating endocytosis machineries.

PAK1 is a serine/threonine kinase found in the cytoplasm and nucleus of cells and PAK1 is important in regulating cytoskeleton remodelling, phenotypic signalling, gene expressions and it affects a variety of cellular processes ^42^. In addition, PAK1 acts downstream of the small GTPases Cdc42 and Rac1, which interact with many effector proteins, including Arp2/3, which in turn can have an effect on cytoskeleton reorganization ^53^. In addition, the role of CDC42 and Rho-GTPases in TNT formation is not fully investigated. A report ^54^, demonstrated that the activity of CDC42 and Rho-GTPases positively contributes to the formation of TNTs in macrophages. In contrast, another study ^55^, found that CDC42/IRSp53/VASP negatively regulates the formation of TNTs. PAK activation can occur independent of Rac and CDC42, the specific lipids particularly sphingolipids can directly activate PAK ^56^. Aβ internalization in primary neurons in absence of apolipoprotein-E has been reported by a cholesterol and sphingolipid sensitive lipids rafts mediated clathrin and caveolae independent but dynamin dependent pathway ^57, 58^. The mechanism is similar to actin-dependent IL2Rβ endocytosis ^40^. AD is highly associated with changes in lipid composition of neurons, and Aβ42 directly downregulates sphingomyelin levels and the ratio of Aβ40/Aβ42 regulates membrane cholesterol ^59^. Here we have detected substantial changes in membrane fluidity upon oAβ treatment using the membrane dye TMA-DPH together with rapid endocytosis. On the other hand, oAβ is implicated in PAK1 dependent synaptic dysfunctions and PAK1 is aberrantly activated and translocated from the cytosol to the membrane during the development of pathology in the AD brain ^60^. Interestingly, HIV and HSV viruses exploit PAK1 dependent endocytosis *en route* and recent studies have also shown direct cell-to-cell spreading of HIV and HSV via TNTs ^19^. Another study has shown that the HIV-1 Nef protein mediated TNT formation is associated with 5 proteins of the exocyst complex and that they are involved in a PAK2 and Rab 11 dependent pathway ^61^. Additionally, alpha herpes virus induced TNT-like membrane actin projections depends on the conserved viral US3 serine/threonine protein kinase-dependent modulations of the cytoskeleton. These modulations are caused by activation of PAK1-dependent signalling and inhibition by IPA-3 attenuates these TNT-like projections ^62, 63^. Exocytosis is involved in the expansion of cell surface area and results in decreased membrane stress. Reduced membrane stress also arises during PM repair, due to extensive exocytosis events ^64^. Endosome recycling also plays a big role in maintaining membrane surface area in equilibrium ^65^. Interestingly, the role of vesicle recycling in TNT formation has been evaluated in a recent study carried out on CAD cells, where an increased level of Rab 11a, Rab 8a and VAMP3 were reported in correlation to TNTs ^66^.

Aβ induced formation of TNTs in primary neurons and astrocytes has been reported earlier ^8^. Here, we have shown the oAβ induced formation of TNTs can transfer aggregates directly between differentiated neurons and the transferred oligomers develop gradual pathology. TNTs as a mean of direct neuron-to-neuron transfer is a convincing model of how oligomers that are suggested as an initiator or driver of AD pathology could gradually progress through the anatomically connected brain regions. However, the possibility of pathology transfer via exosomes could be a parallel mechanism, since we have observed that the formation of TNTs is followed by oAβ induced enhanced lysosomal exocytosis. Therefore, further in-depth studies are needed to understand how cells maintain homeostasis of intercellular communication by balancing exosome release and TNTs in the stressed cells. Altogether, our results have indicated that TNTs are characteristic membrane actin conduit that transfer oAβ aggregates from one cell-to-another, and are formed as a consequence of oAβ induced membrane damage and Ca^2+^ dependent lysosomal-exocytosis engaged rapid membrane repair process, followed by PAK1-kinase dependent CIE and actin remodelling (Fig. 7), probably to maintain cell surface area expansion and membrane stress in equilibrium.

**Figure 7:**
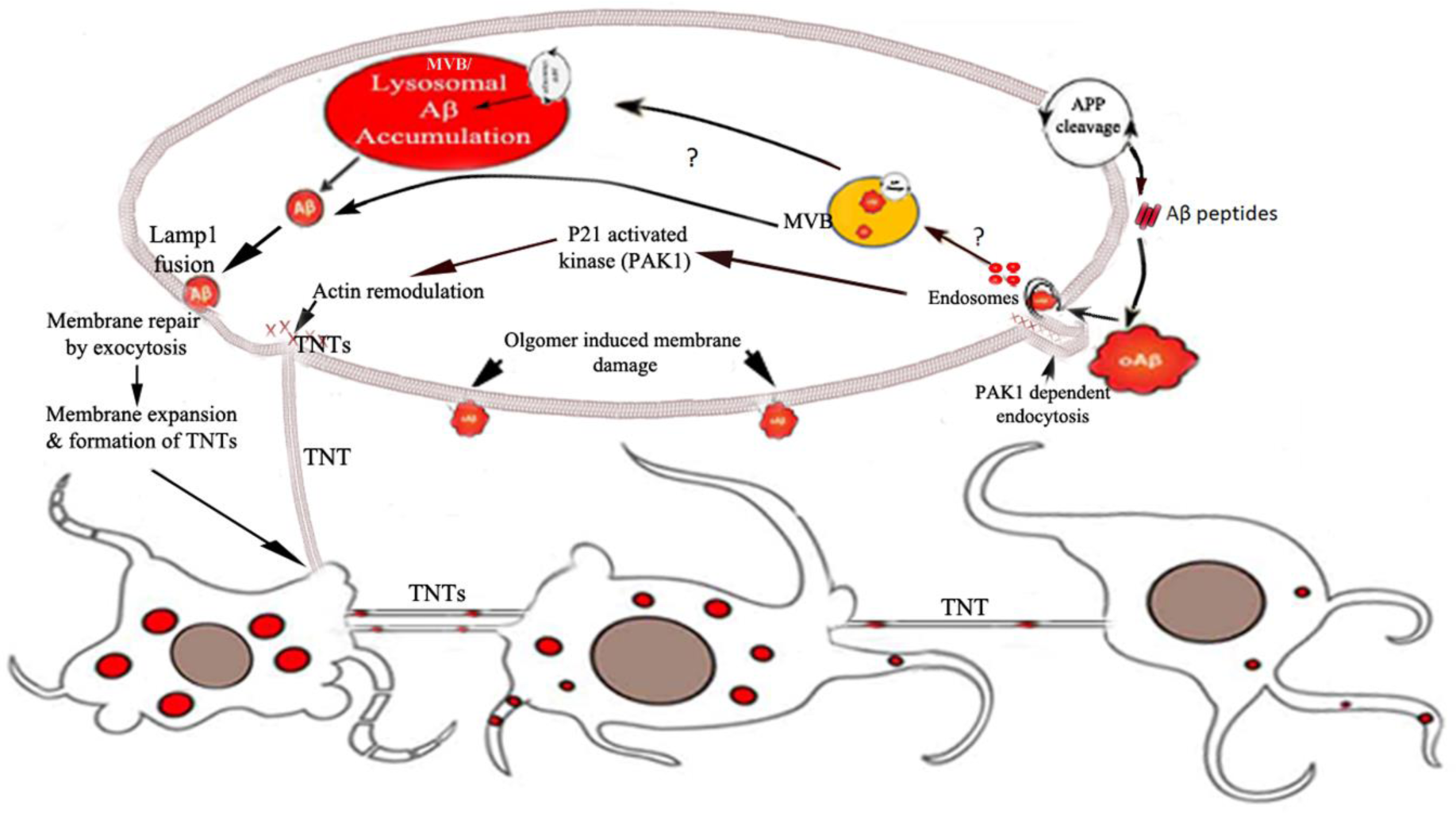
Schematic summary to show that the sprouting of TNTs might be derived as a result of oAβ induced PM damage and Ca^2+^ dependent lysosomal-exocytosis engaged rapid membrane repair process, followed by PAK1-kinase dependent CIE and actin remodelling to re-establish the cell surface area expansion or membrane stress in equilibrium.

## Acknowledgement

SN thanks to Manipal Academy of Higher Education intramural grants (India; 2019-2022), Magnus Bergvalls (Sweden, 2015-2016, #2014-00192), Gun och Bertil Stohnes research grants (Sweden, 2014-2015) and Alzheimer fonden (Sweden; 2012-2013). We thank our long-term collaborator Prof. Martin Hallbeck of Linkoping University, Sweden, for the valuable suggestions on designing of cell-to-cell transfer experiments. We thank Dr. Ravi Manjithaya of JNCASR, Bangalore, India, for giving us SH-SY5Y cell lines and giving access to the SIM-superresolution microscope. Thanks a lot to Likhesh Sharma, PhD, Product Manager GE Healthcare for assisting with imaging by SIM-superresolution microscope. We thank Prof. Satyajit Mayor of National centre for biological sciences, India, for his kind gift of GFP-GPI construct, CAAX-mCherry plasmid, sharing of some reagents and for his valuable inputs. The CAAX-mCherry plasmid was a kind gift to Prof. Satyajit Mayor by Prof. Jacco van Rheenen of Hubrecht Institute for Developmental Biology and Stem Cell Research. Thanks to Prof. Subba Rao Gangi Setty of Indian Institute of Science and Prof. Gopal Pande of Manipal institute of higher education, India, for their valuable inputs.

## Author Contributions

S.N conceived and conducted the research; S.N and K.O designed research; S.N designed tunneling nanotubes, oAβ internalization and PAK1 experiments; K.O. designed membrane dynamics and membrane repair experiments; A.D, D.K.V, N.D, C.K and S.N performed experiments and analysed the data; S.N, K.O and K.K interpreted data; S.N wrote the paper taking valuable inputs from all the authors.

Authors have no Competing Interests.

