## Supplementary Material for "Amyloid-β induced membrane damage instigates tunneling nanotubes by exploiting p21-activated kinase dependent actin remodulation" (from another version of this preprint: version 3, 2021-06-13)

[3] Current address: Novo Nordisk Foundation Center for Protein Research, University of Copenhagen, Copenhagen, Denmark.

+ Equally contributed

**Supplementary Figures and Figure legends**

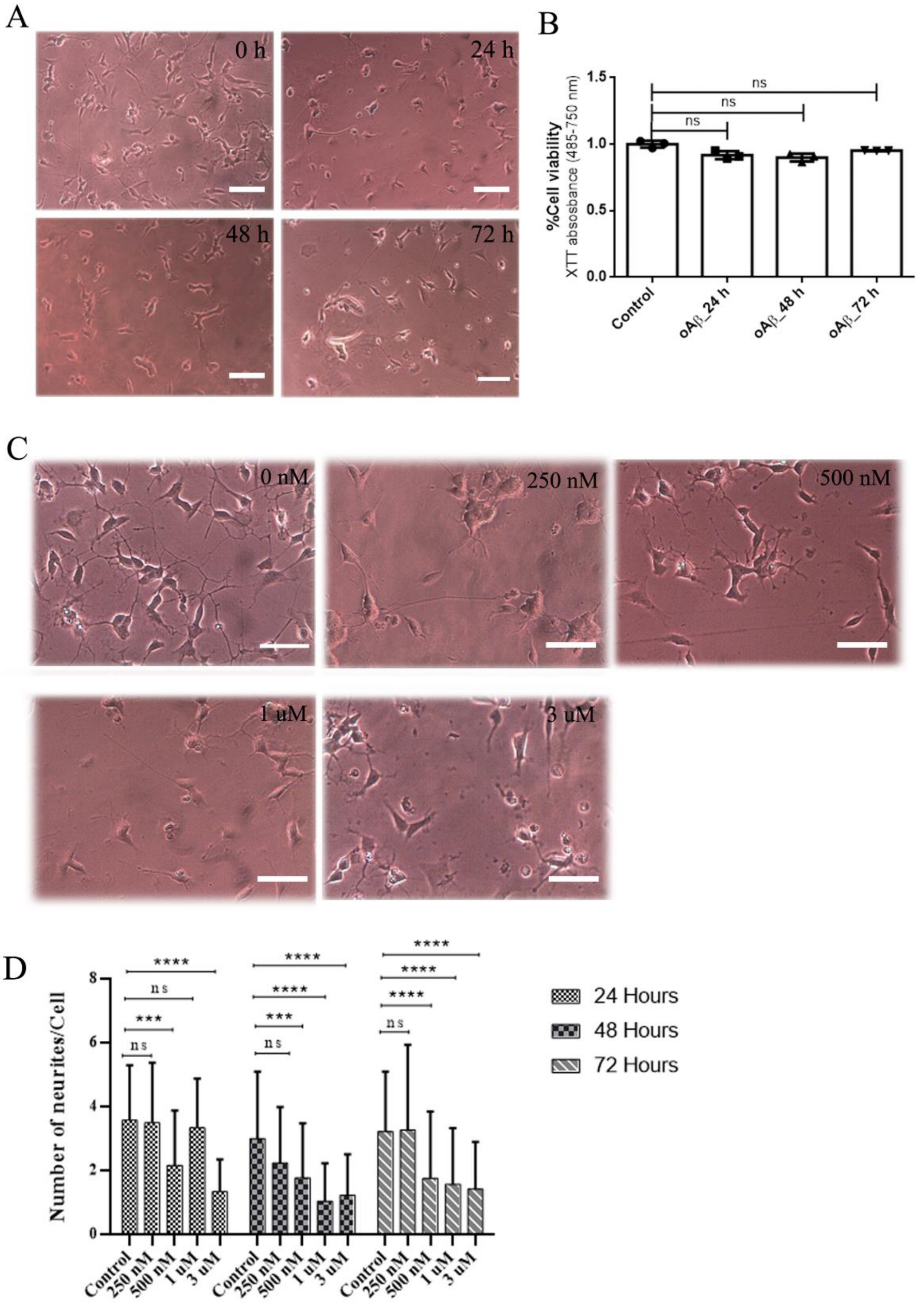


**Supplementary Figure 1. Time dependent and concentration dependent cytotoxicity effects of oAβ on differentiated cells.** A) Time dependent cytotoxicity effects of oAβ (1µM) on differentiated cells, imaged by bright field mode. Cells show fewer numbers of neurites over the time of oAβ treatment. B) However, cells show no significant change in cytotoxicity measurements, while quantified by XTT assays. One-way ANNOVA test were performed. C) Concentration dependent cytotoxicity effects of oAβ at 24 h on differentiated cells. D) Cells show fewer numbers of neurites with increasing concentration of oAβ treatment. Two-way ANNOVA test were performed to validate statistical significance. Scale bars are 50 µm.

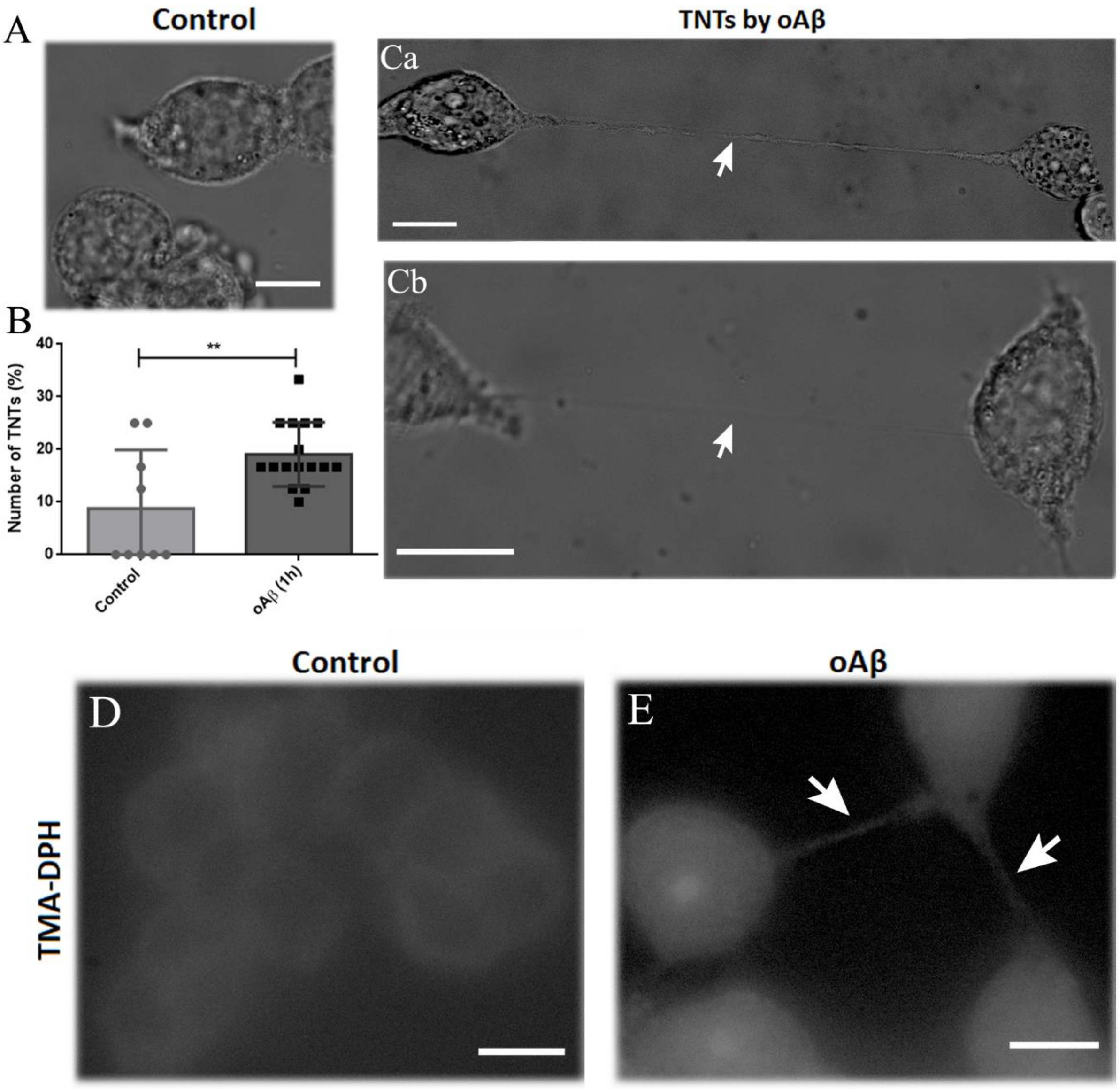


**Supplementary Figure 2. oAβ can induce TNTs in non-neuronal cells.** A) Control HEK293 cells. B) TNT- like structures were counted and plotted in percentage with respect to the number of cells from each of the image frames and numbers of cells > 60 for each sets. C) Formation of TNT between two cells in oAβ (1 µM for 1 h) treated cells (indicated by white arrows). Parts of sliced images were stitched (a: 3 slices and b: 2 slices) to observe the long TNTs as whole. D-E) HEK293 cells were incubated with 1 µM of oAβ for 1 h together with a membrane impermeable dye TMA-DPH. D) In control cells, the TMA-DPH shows labelling of intact peripheral PM. E) In contrast, in oAβ treated cells formation of TNT-like structures (indicated by white arrows) and there is a change in membrane fluidity, which causes the permeabilization of the dye TMA-DPH. Scale bars = 10 µm.

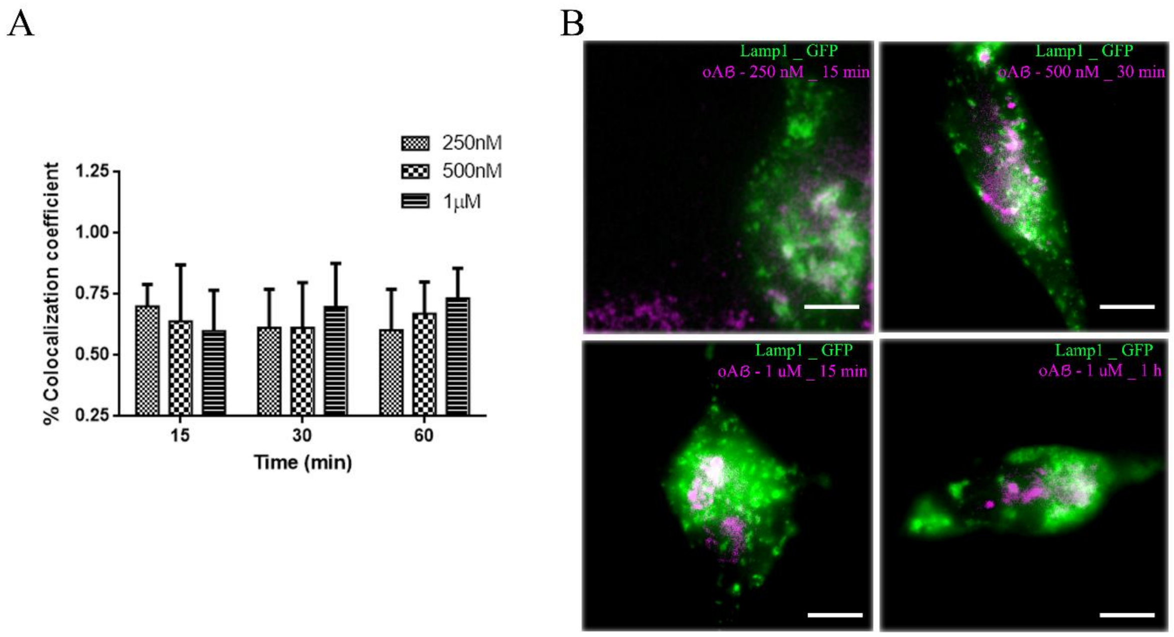


**Supplementary Figure 3.** A) Internalization of extracellularly applied oAβ in the SHSY-5Y cells at various concentrations (250 nM to 1 µM) and over the time of incubations (15 min to 1 h) into endo-lysosomal pathways were quantified by analyzing colocalization with LAMP1 protein, the marker of late endosomes or multi vesicular bodies (MVB). B) Representative images are showing significant colocalization of the oAβ-TMR (magenta) in the LAMP1-GFP(green) positive vesicles at different concentrations and for different periods of incubations time. Scale bar=10µm.

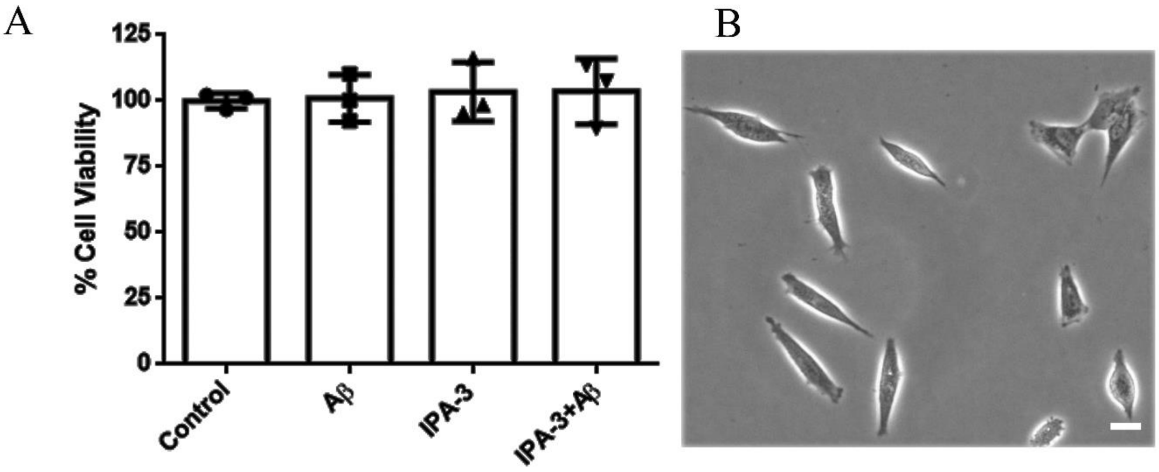


**Supplementary figure 4.** A) IPA-3 (20 µM) pretreatments (30 min) with and without oAβ (1 µM; for 1 h) co-treatment show no change in cell viability, while quantified by MTT assays. B) Phase contrast image of undifferentiated SH-SY5Y cells. Sparsely distributed undifferentiated cells were used to identify and count properly the TNTs between relatively distant cells. Scale bar = 10 µm.

**Figure legends of Supplementary Movies.**

**Supplementary Movie 1.** Image sequences to show clearly the connected nanotubes (white arrow) between neighbouring cells detected in differentiated SH-SY5Y cells incubated for 3 h with 500 nM of oAβ-TMR (magenta), washed and stained with 50 nM of lysotracker (green) before capture of the video. The cells with long TNTs form noticeable blebs (yellow arrow) and cell membrane expansion. From the video one can find out that membrane nanotube (white arrow) stays as hung conduit like structure between the two connected cells without touching the surface of the culture dish.

**Supplementary Movie 2.** TNT formation depends on the concentration of oAβ. Degradation of neurites are not significant at 100 nM of oAβ (unlabelled) incubated for 3 h (200 nM of lysotracker green was added before imaging), instead, formation of few TNT like structures (white arrows) extended from lamellipodia are distinctly visible at this condition. Membrane of the cells are often expanded with blebs or lamellipodia and organelles like granular structures were noticeable to travel in the TNTs (blue arrow).

**Supplementary Movie 3.** We have also observed propagations of organelles (blue arrows) via TNTs (white arrows) in differentiated cells (yellow arrows) treated with 250 nM of oAβ-TMR (magenta) for 3 h. At 250 nM, the healthy neurites are fewer in numbers and membrane of the cells are often expanded with blebs or lamellipodia (yellow arrows) and oligomers of oAβ-TMR in granular structures like organelles (magenta) were noticeable to travel in the TNTs (blue arrows).

**Supplementary Movie 4.** We have followed the transfer of oAβ-TMR via TNTs from one cell-to-another in differentiated cells by time-lapse imaging. Cells were incubated with 500 nM of oAβ-TMR (magenta) for 3 h, washed and labelled with 50 nM of lysotracker (green) and time lapse image sequences were taken. The cells form TNTs and oAβ-TMR travels from one cell-to-another as organelles like puncta structures through the TNTs (blue arrows).

**Supplementary Movie 5.** Control undifferentiated SH-SY5Y cells with the membrane dye TMA-DPH (0.5 µM). Control cells show no blebs, filopodium or TNT structures.

**Supplementary Movie 6.** Undifferentiated SH-SY5Y cells incubated with 1 µM of oAβ together with the membrane dye TMA-DPH (0.5 µM) and imaged immediately after addition. TNTs formation was observed within 10 min, preceding of massive membrane activities, filopodium formation and endocytosis of membranes.
